# Genetic and environmental risk for chronic pain and the contribution of risk variants for psychiatric disorders. Results from Generation Scotland: Scottish Family Health Study and UK Biobank

**DOI:** 10.1101/037457

**Authors:** AM McIntosh, LS Hall, Y Zeng, MJ Adams, J Gibson, E Wigmore, AM Pujils-Fernandez, AI Campbell, T-K Clarke, C Hayward, C Haley, DJ Porteous, IJ Deary, DJ Smith, BI Nicholl, DA Hinds, AV Jones, S Scollen, W Meng, BH Smith, LJ Hocking

**Affiliations:** Division of Psychiatry, University of Edinburgh, Royal Edinburgh Hospital, Edinburgh EH105HF; Centre for Cognitive Ageing and Cognitive Epidemiology, University of Edinburgh, 7 George Square, Edinburgh EH8 9JZ; Institute for Genetics and Molecular Medicine, University of Edinburgh, Western General Hospital, Crewe Road, Edinburgh EH4 2XU; Institute of Health and Wellbeing, University of Glasgow, Glasgow G12 8RZ; 23andMe Inc., Mountain View, California, United States of America; Pfizer WRD, Human Genetics and Computational Biomedicine, The Portway Building, Granta Park, Cambridge CB21 6GS; Division of Population Health Sciences, University of Dundee, Ninewells Hospital and Medical School, Kirsty Semple Way, Dundee DD2 4BF; The Institute of Medical Sciences, University of Aberdeen, Forseterhill, Aberdeen AB25 2ZD

## Abstract

**Background:** Chronic pain is highly prevalent worldwide and a significant source of disability, yet its genetic and environmental risk factors are poorly understood. Its relationship with psychiatric illness, and major depressive disorder (MDD) in particular, is of particular importance. We sought to test the contribution of genetic factors and shared and unique environment to risk of chronic pain and its correlation with MDD in Generation Scotland: Scottish Family Health Study (GS:SFHS). We then sought to replicate any significant findings in the UK Biobank study.

**Methods:** Using family-based mixed-model analyses, we examined the contribution of genetics and environment to chronic pain using spouse, sibling and household groups as measures of shared environment. We then examined the correlation between chronic pain and MDD and estimated the contribution of genetic factors and shared environment. Finally, we used data from two independent genome-wide association studies to test whether chronic pain has a polygenic risk architecture and examine whether genomic risk of psychiatric disorder predicted chronic pain and whether genomic risk of chronic pain predicted MDD.

**Results:** Chronic pain is a moderately heritable trait (narrow sense heritability = 38.4%) which is more likely to be concordant in spouses and partners (variance explained 18.7%). Chronic pain is positively correlated with depression (rho = 0.13, p = 2.72x10^−68^) and it shows a tendency to cluster within families for genetic reasons (genetic correlation rho = 0.51, p = 8.24x10^−19^). Polygenic risk profiles for pain, generated using independent GWAS data, predicted chronic pain in both GS:SFHS (maximum β = 6.18x10^−2^, p = 4.3x10^−4^) and UK Biobank (maximum β = 5.68 x 10^−2^, p < 3x10^−4^). Genomic risk of MDD is also significantly associated with chronic pain in both GS:SFHS (maximum β = 6.62x10^−2^, p = 4.3x10^−4^) and UK Biobank (maximum β = 2.56x10-2, p < 3x10^−4^).

**Conclusions:** Genetic factors and chronic pain in a partner or spouse contribute substantially to the risk of chronic pain in the general population. Chronic pain is genetically correlated with MDD, has a polygenic architecture and is predicted by polygenic risk of MDD.

## Introduction

Chronic pain and major depressive disorder (MDD) are highly prevalent and frequently comorbid conditions [1] that complicate chronic physical disease and are leading causes of disability globally in the general population (Lancet 386:743-800). The causes of chronic pain and depression are poorly understood, although there is some evidence of a contribution from genetic factors in each disorder [2, 3].

Using a family-based approach involving 7644 individuals from 2195 extended families from Generation Scotland, Hocking et al demonstrated that severe chronic pain has a heritability of 30% [4]. In twin studies the broad sense heritability of chronic pain has also been estimated similarly at 32% [5]. The genetic architecture of chronic pain is however unknown and the potentially polygenic contribution of many common variants to differences in liability is, to date, unproven. The heritability of MDD has been estimated, from a metaanalysis of twin studies, at around 37% [6, 7], similar to estimates for chronic pain. Whilst the majority of the heritability of MDD is not explained by the risk variants discovered to date, the genetic architecture of MDD is understood to be highly polygenic, with multiple loci of low-penetrance acting additively to confer liability to the disorder [8, 9].

The phenotypic associations of both chronic pain and depression are also numerous and overlapping [10-15]. Since chronic pain and depression are amongst the most disabling conditions worldwide, it is important to understand the nature of their co-occurrence and whether this is caused by shared genetic or environmental factors. Studies that quantify the relative roles of genetics, and shared and unique environment in chronic pain, including those that seek to explain its correlation with depression, are small in sample size and number. One study of 3266 participants/twins estimated the cross-trait correlation between depression and pain, and found a latent trait explained the covariance between chronic widespread pain and major depression that had 86% heritability [16]. Such studies may focus attention on avenues for future therapeutic intervention, help to clarify how these two common conditions are related and provide clues to how risk may be mitigated at a population level.

The main aim of the current study was to assess the genetic and environmental contributions to pain and its comorbidity with MDD. Using data from Generation Scotland: Scottish Family Health Study (GS:SFHS) [17], we estimated the magnitude of the contribution to chronic pain from genetic factors, and from shared and unique environmental influences using pedigree-based analyses. Next, we measured the correlation between chronic pain and MDD using bivariate pedigree-based analysis. We then conducted a polygenic profile scoring analysis of GS and UK Biobank using independent data available from two large-scale genome-wide association studies (GWAS) of chronic pain and MDD. In doing so, we first tested whether there was evidence of a polygenic architecture for pain. Secondly, we examined whether polygenic risk of MDD predicted chronic pain using molecular genetic data.

## Methods

Two cohorts were used to examine the genetic and environmental contributions to chronic pain and its relationship with depression: GS:SFHS (N=23,960) and UK Biobank (UKB, N=112,151). GS:SFHS, having a family-based design, was used to estimate the contributions of genetic factors, and of shared and unique environment to chronic pain and its correlation with MDD, psychological distress and neuroticism. GS:SFHS and UKB were then both used to test the genetic relationship between chronic pain and depression risk scores and their associated phenotypes using polygenic risk score analysis. Marker effects were estimated from independent GWAS studies that did not include either the GS:SFHS or UKB study samples.

### Generation Scotland: Scottish Family Health Study

GS:SFHS is a family-structured, population-based cohort recruited at random through general medical practices across Scotland. The protocol for recruitment is described in detail elsewhere [17, 18]. The cohort consists of 23,960 individuals over 18 years of age who were recruited if they had at least one other family member willing to participate. Pedigree information was available for all participants, and this has subsequently been validated against estimates of relatedness estimated using genome-wide single nucleotide polymorphism (SNP) data on 19,995 individuals. All individuals from GS:SFHS provided data for the current analyses (median age = 47.9, 41% male, 59% female) of whom 19,511 also provided genotype data for risk profiling after quality control. (mean age = 47.50, SD = 14.98) (N = 11,514 female, N = 7,997 male).

Using pedigree information in GS:SFHS, we have created variables to represent the random effect from genetic and environment factors for linear mixed modelling. We created the genetic relationship matrix using all participants as a means of estimating the heritability of specific traits and their genetic correlation between trait pairs. In addition, we fitted several variables in the model to estimate the effects of shared environment. These included measures of common household, common spouse/partner and a measure representing siblings that shared a common parental and sibling environment. Further details of these measures are provided in the supplementary material.

### Assessment of chronic pain in Generation Scotland

GS:SFHS participants (N = 16955) completed a questionnaire assessment of chronic pain as previously described by Hocking [19]. This included, first, a validated chronic pain identification questionnaire, in which chronic pain was ascertained in the event of positive responses to two questions: (1) Are you currently troubled by pain or discomfort, either all the time or on and off? and (2) Have you had this pain or discomfort for more than 3 months [20, 21]. Secondly, severity of chronic pain was determined by the Chronic Pain Grade (CPG) questionnaire, a validated 7-item instrument that measures the pain intensity and pain-related disability and grades severity from Grade 1 (low intensity, low disability chronic pain) to Grade 4 (high intensity, high disability, severely limiting chronic pain)[21-23].

### Assessments of MDD in Generation Scotland

All GS:SFHS participants were invited to complete a screening questionnaire for MDD as described previously [24]. Those screening positive were invited to complete the Structured Clinical Interview for the Diagnostic and Statistical Manual of the American Psychiatric Association (SCID, DSM version 4) and were subsequently diagnosed with MDD if they fulfilled DSM-IV criteria. Individuals who screened negative for MDD or who screened positive but did not fulfil criteria were diagnosed as being free from MDD. Individuals who declined to complete the screening questionnaire or the SCID had their MDD status set to missing. Details of the genotyping and quality control procedures are given in the supplementary material.

All GS:SFHS individuals provided written informed consent to participation and for use of their data and samples for medical research. Ethical approval for the study was given by the NHS Tayside committee on research ethics (reference 05/s1401/89).

### UK Biobank

UK Biobank is a health research resource that aims to improve the prevention, diagnosis and treatment of a wide range of illnesses. Between the years 2006 and 2010, approximately 500,000 people were recruited from across the UK [25]. For the present study, 112,151 community-dwelling individuals (58,914 females, 53,237 males) aged 40 to 73 years (mean = 56.91 years, SD = 7.93) with genome-wide genotyping were available. Individuals were assigned to self-declared MDD if they stated they had previously suffered from clinical depression in the past during their nurse-led assessment (variable no 20002).

### Assessment of chronic pain in UK Biobank

To identify chronic pain participants were asked, ‘In the last month have you experienced any of the following that interfered with your usual activities?’: headache, facial pain, neck or shoulder pain, back pain, stomach or abdominal pain, hip pain, knee pain, pain all over the body. With each positive response, they were asked whether the pain had lasted for at least 3 months, and individuals who reported at least one of these pains lasting for at least 3 months were defined as having ‘chronic pain’. Chronic pain in UK Biobank was then quantified based on a previous report [1] which categorised chronic pain in ascending order based on whether a single site was affected, 2-3 sites were affected or if the pain was widespread and diffuse, affecting 4-7 sites or “pain all over the body”.

UK Biobank (Collins 2012, http://www.ukbiobank.ac.uk) received ethical approval from the Research Ethics Committee (reference 11/NW/0382) and the present analyses were conducted under UK Biobank data application numbers 4844 and 3501.

### Polygenic profiling

Genetic risk of pain was estimated using an independent data set designed by Pfizer conducted in collaboration with 23andMe on their participants. In this Pfizer-23andMe study, validated pain questionnaires, using identical case and severity measures to those used in GS:SFHS, were completed by >32,000 research participants. 23andMe research participants provided informed consent to take part in this research under a protocol approved by the Ethical and Independent Review Services, an AAHRPP-accredited institutional review board. The number of unrelated Europeans that were included after relatedness removal was 10,780 cases and 12,552 individuals categorised as ‘no pain’ controls. Chronic pain phenotypes for 23andMe participants were defined (as for GS:SFHS). For each pain phenotype, individual SNP effects (odds ratios and 95%Cls) were calculated in PLINK using a linear regression analysis after adjustment for age, sex, body mass index (BMI), current manual labour and previous manual labour and the first five population principal components. Further details of the genotyping and quality control performed by 23andMe are provided in the supplementary material.

Genetic risk of MDD was estimated using a previously published independent GWAS data sets from the Psychiatric Genomics Consortium (PGC) Major Depressive Disorder Working Group. Details of the quality control, imputation and analysis have been previously provided in a published manuscript [8].

Chronic pain and MDD polygenic risk profile scores for all individuals in GS:SFHS and UKB with useable GWAS data were then estimated. Polygenic risk profiles were estimated genome-wide using four arbitrarily defined nominal p-value thresholds in the original discovery GWAS (p-values = 0.01, 0.05, 0.1 and 0.5) producing a score in a second dataset at each corresponding threshold. Statistically significant prediction of chronic pain in GS:SFHS and UKB using marker weights estimated in Pfizer-23andMe GWAS data was assumed to infer a polygenic risk architecture. Statistically significant prediction of chronic pain using MDD polygenic profile scores (or vice versa) was assumed to infer an overlapping polygenic risk architecture.

## Statistical analysis

### Pedigree-based analysis of chronic pain

Pedigree-based analysis of chronic pain was conducted solely in the GS:SFHS dataset using the ‘Chronic Pain Grade’ (CPG 0-4) variable. Using a generalised linear mixed-model, we estimated the relative contributions of the random effect from (1) genetic factors (using the additive genetics (or numerator) relationship matrix) calculated from the pedigree, and (2) environmental factors from the parent (using the sib variable), previous/current spouse (using the spouse variable) and household (using the two household variables). The magnitude of each environmental effect (parent, spouse, household) was estimated in the presence of a genetic variance component to reduce confounding of each environmental effect by genetic relatedness. All fixed and random effects were estimated in the MCMCglmm package for Bayesian Generalised linear mixed models (Hadfield Journal of Statistical Software 2010) and in the R package ASReml-R [26] (version 3) in R version 3.2.2. Since MCMCglmm can adequately model the genetic and environmental contributions to binary and ordinal traits using Markov Chain Monte Carlo sampling, MCMCglmm was used as the primary methodology for estimating the effects sizes for each variance component. Sex, age and age^2^ were used as fixed effects throughout.

In order to select the best-fitting model to the data, we employed two convergence methodologies in each software package. Firstly, we estimated goodness-of-fit in the MCMCglmm package using Deviance Information Criterion (DIC) with larger values indicating poorer fitting models. Secondly, we used ASReml-R to compare goodness of fit by estimating the difference in log-likelihood between two models, which was then compared to a chi-squared distribution with the relevant degrees of freedom for the number of parameters that differed between the models. A significant p-value indicated a better fitting model with a lower log-likelihood. In each case we started with a model containing only a genetic effect, then applied models containing the genetic effects plus one of the three environmental effects (parent, spouse and household). After choosing the best model from those estimated, we attempted to improve the model fit by adding in a second environmental variable. Where the DIC or log-likelihood could not be further improved, we selected the reduced model as the best fit for the available data and provided the effects from MCMCglmm.

### Pedigree-based bivariate analyses of pain and MDD

In order to examine the relationship between chronic pain and MDD in GS:SFHS, we used a generalised linear mixed model as implemented in ASReml-R. We estimated the phenotypic correlation between chronic pain and MDD first before providing the genetic correlation coefficient and environmental coefficient for any significant shared environments estimated from the univariate model. The significance of each correlation was estimated using two nested ‘null’ models. The first ‘null’ genetic model was constrained to have a genetic correlation of zero, whereas the ‘null’ environmental model was constrained to have an environmental correlation of zero. Each model was compared to the full (estimated) model and the significance of each effect was determined using the likelihood ratio test (LRT) by comparing 2 times the difference in the log likelihood to a chi-squared distribution with 1 degree of freedom. Standard errors for each correlation coefficient were also determined using the variance components provided by ASReml-R. In order to account for relatedness and to adjust for the fixed effects of sex, age and age^2^, the significance of the phenotypic correlation coefficient was estimated using the ratio of its effect size to its standard error in ASReml-R. The significance was then estimated by comparison of this figure to a z-distribution and the calculation of a two-tailed p-value.

### Polygenic profiling

Polygenic profiling of the GS:SFHS sample was conducted using the marker weights estimated in the Pfizer-23andMe dataset for the CPG trait in this independent discovery sample. These were first used as predictors of chronic pain in GS:SFHS using the profile score as a fixed effect within a generalised linear mixed model implemented in MCMCglmm. Additionally, sex, age, age^2^ and four multidimensional scaling components were also used as fixed effects. Each model was adjusted to take account of relatedness between subjects using the relationship matrix. The significance of the polygenic risk profile effect in predicting chronic pain in GS:SFHS was estimated using the modal value and the 95% credible interval from the posterior density. The variance explained by the polygenic profile score was estimated by multiplying the profile score by its corresponding regression coefficient and estimating its variance ([27] in ASReml-R. This value was then divided by the variance of the observed phenotype itself to yield a coefficient of determination between 0 and 1. Polygenic profile scores for chronic pain were then used to predict MDD in GS:SFHS.

We then profiled all individuals in GS:SFHS using the publically available PGC dataset for MDD. Polygenic profile scores for MDD were then tested for their association with MDD in GS:SFHS as a means of assessing their validity and to identify a theoretical upper bound for their accuracy in predicting chronic pain. Secondly, we estimated the strength of the association between polygenic profile score for MDD and chronic pain in GS:SFHS using the same models as described in the previous paragraph.

Where a significant association was found between polygenic risk of chronic pain and either chronic pain or MDD in GS:SFHS, we repeated the analysis in UKB using their different chronic pain phenotype which is defined, in contrast to GS:SFHS, on the extent rather than on the intensity of the chronic pain. Where significant associations were found between polygenic risk of MDD and chronic pain in GS:SFHS, these analyses were repeated excluding individuals with major depression (experienced at any point in their life) in order to test whether the same relationship between polygenic risk of MDD and chronic pain was present in the unaffected population.

Associations that were shown to be significant in GS:SFHS were then repeated in UKB using unrelated individuals only. Polygenic risk profile scores were examined for their association with observed phenotypes in ASReml-R using the same methods, but without the inclusion of a GRM due to the large data set and unrelated nature of the filtered UKB study population used in the current investigation. Further details of UKB are provided in the supplementary materials.

## Results

### Univariate analysis of chronic pain grade in GS:SFHS

Five models were initially estimated for the chronic pain (CPG) dependent variable. In each case the model included a genetic effect, then each of the three environmental effects individually (‘sib’ (common parent), spouse or household). The model including the additive genetic effect plus that of spouse was a better fit than the model that only included the additive genetic effect alone either by selection of the model with the lowest DIC, or through the likelihood ratio test (LRT). The effect of additive genetic effects and spouse alone was also superior (in terms of DIC) to either of the other two models that included additive genetic effects and ‘sib’ (common parent) alone, or additive genetic effects and household alone. In a second step, we added further environmental effects to the model that included additive genetics and spouse alone, but we could not improve on the model fit adjudicated using either the DIC or LRT. All of the models are shown in Supplementary Table 1 within the Supplementary Material.

**Table 1:**
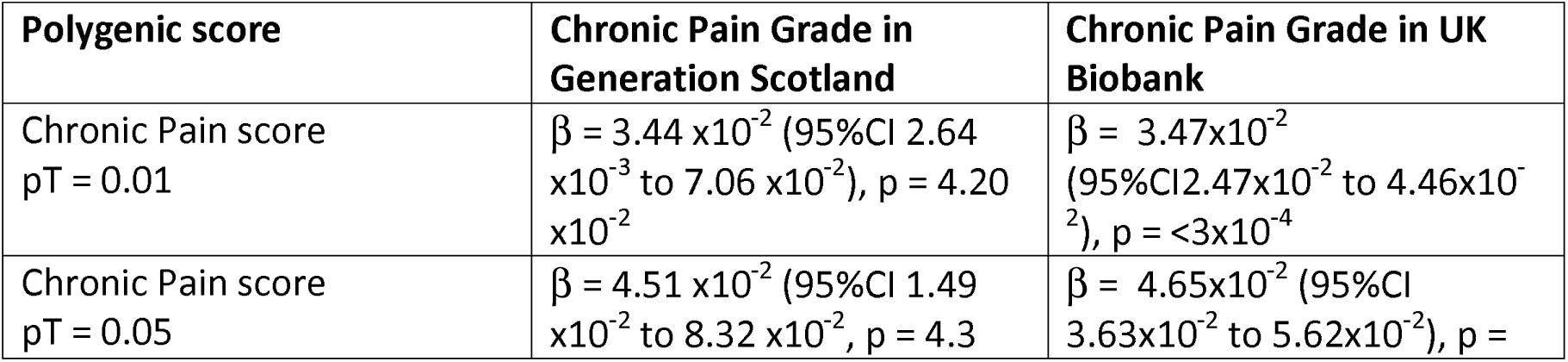
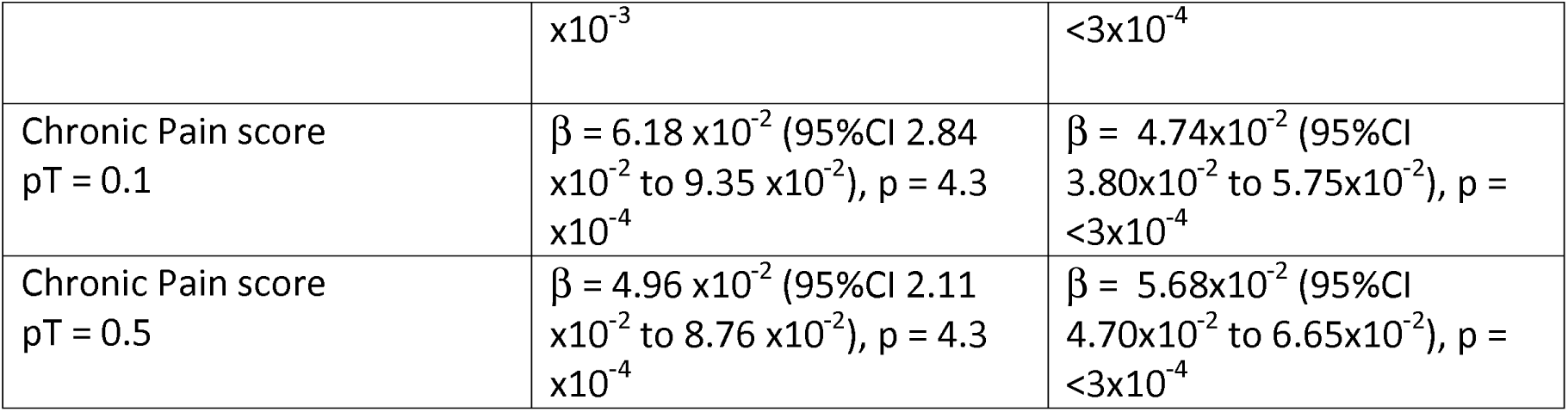
Association between Pfizer-23andMe derived polygenic profiles scores for chronic pain and chronic pain in GS:SFHS and UK Biobank. Beta: represents standardised regression coefficient between the four polygenic risk scores and the two chronic pain phenotypes. All analyses were conducted in MCMCglmm. The 95% CI represents the 95% credible interval, which is broadly interpreted as the interval in which there is a 95% probability that the true parameter lies.

The heritability of chronic pain was substantial, accounting for 38.4% (95%CI 33.6% to 43.9%) of the variation in CPG score. Shared environment with a spouse accounted for approximately 18.7% (95%CI 9.5% to 25.1%) the variation in liability to chronic pain. The magnitude of this effect was approximately half of the variance component explained by additive genetic factors. The residual unexplained variance, reflecting measurement error, poor reliability and non-shared environment, accounted for just under half of the variance in liability to chronic pain (variance attributed 42.9%).

### Bivariate analysis of chronic pain grade in GS:SFHS

Chronic Pain was positively correlated in GS:SFHS with MDD (Pearson’s phenotypic correlation: r = 0.13, SE = 0.01, p = 2.72x10^−68^). Phenotypic covariance between chronic pain and MDD was demonstrably attributable to shared genetic architecture (Pearson’s r = 0.51, SE = 0.054, p = 8.24x10^19^) and also to a spouse/partner environmental effect (Pearson’s r = 0.53, SE = 0.24, p = 0.02). The larger standard errors (SE = 0.24) for the effect of partner/spouse likely reflected the smaller number of spouse/partners (N = 3,486) and the lower precision in these estimates.

### Effects of polygenic risk of pain on chronic pain in GS:SFHS and UK Biobank

Polygenic risk profile scores for the Pfizer-23andMe phenotype ‘Chronic Pain Grade’ and significantly predicted chronic pain in both the GS:SFHS and UK Biobank samples, at all p-value thresholds (Table 1). In each case, the proportion of variance in chronic pain explained by the polygenic risk score was less than 1%. In contrast, chronic pain polygenic risk profile scores derived from Pfizer-23andMe marker data were not associated with MDD in GS:SFHS at even nominal levels of significance (Table 2).

**Table 2:**
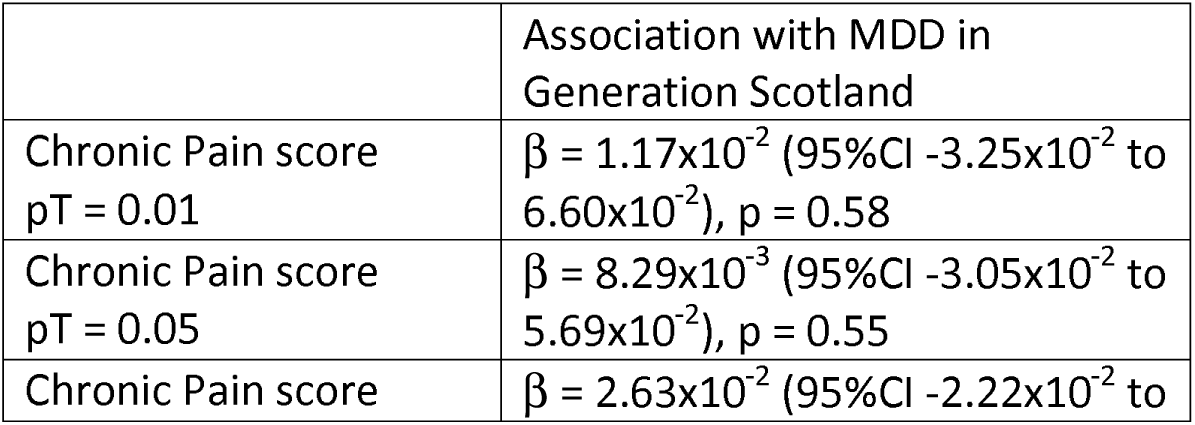
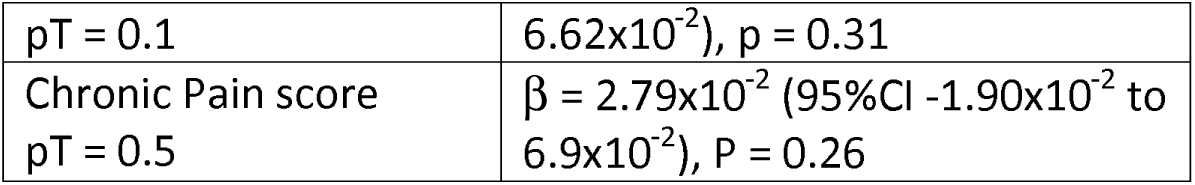
Association of polygenic profile scores for chronic pain and MDD in GS:SFHS. Betas represent the standardised regression coefficient between each pain polygenic risk score and MDD in Generation Scotland (GS:SFHS)

**Figure 1:**
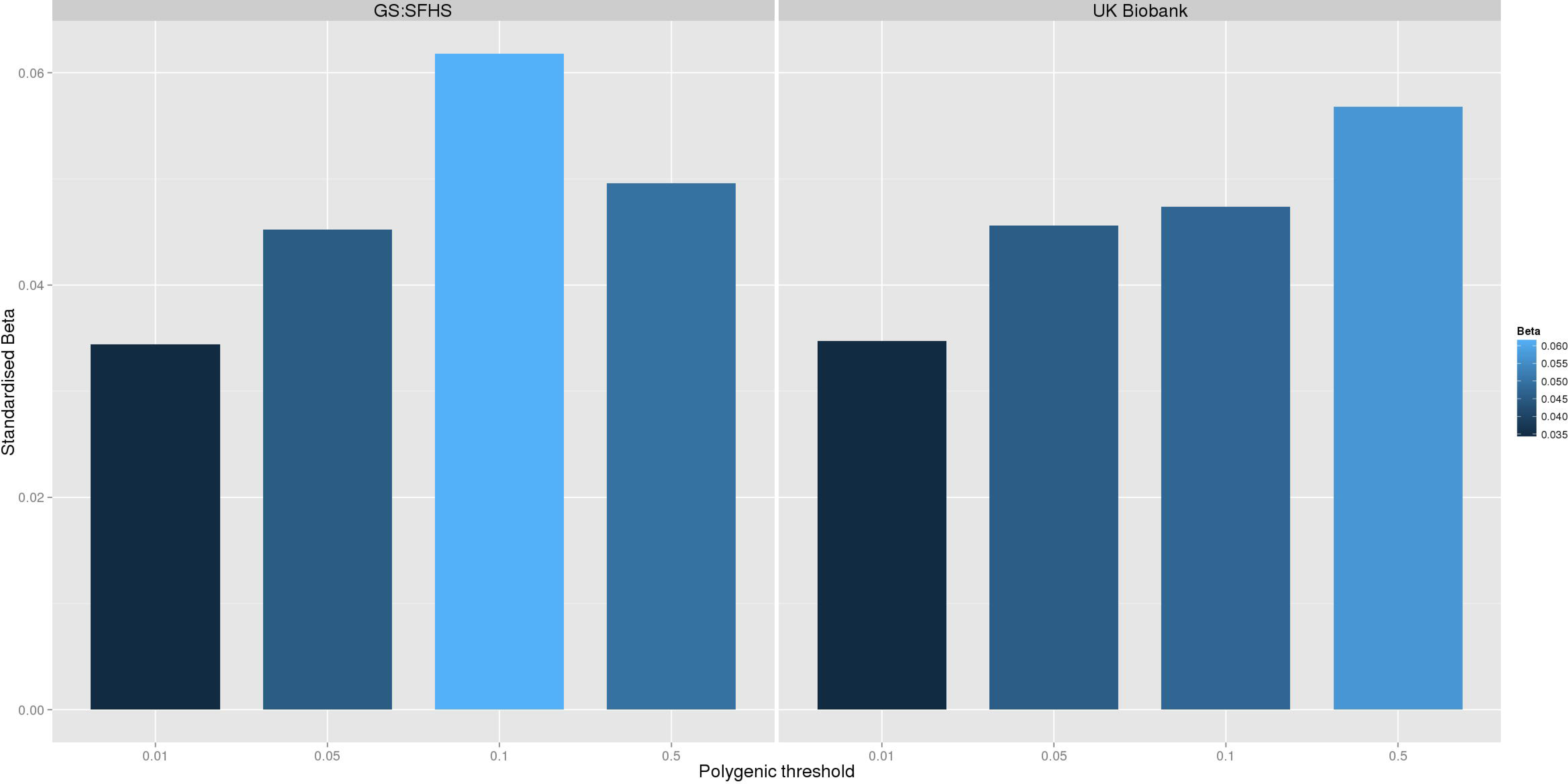
Effect of Pfizer-23andMe derived polygenic risk profiles scores for pain on chronic pain phenotypes in GS:SFHS (left panel) and UK Biobank (right panel) shows the association between polygenic risk scores for pain (derived from Pfizer-23andMe data) and chronic pain in GS:SFHS (left panel) and UK Biobank (right panel). Vertical y-axis represents the effect size as a standardised beta, horizontal axis represents the four alternative p-value thresholds used for the generation of polygenic scores in the discovery GWAS studies.

### Effects of polygenic risk for MDD in GS:SFHS

Polygenic risk profile scores for MDD were estimated in GS:SFHS and UK Biobank using data from the PGC international consortia. Polygenic risk profile scores for MDD were significantly positively associated with MDD in GS:SFHS at 3 out of 4 thresholds, with the exception of the pT = 0.01 threshold. In the larger UK Biobank study, polygenic risk of MDD was associated with MDD at all 4 p-value thresholds (Table 3).

**Table 3:** Prediction of MDD related traits in GS:SFHS from Pfizer-23andMe derived pain scores and Psychiatric Genomics Consortium MDD polygenic risk scores. Beta: represents standardised regression coefficient between the four polygenic risk scores and MDD in GS:SFHS and UK Biobank. All analyses were conducted in MCMCglmm. The 95% CI represents the 95% credible interval, which is broadly interpreted as the interval in which there is a 95% probability that the true parameter lies.

| PGRS score | MDD in GS:SFHS | MDD in UK Biobank |
| --- | --- | --- |
| MDD score<br>pT = 0.01 | $\beta = 3.18 \times 10^{-2}$ (95%CI $-1.56 \times 10^{-2}$ to $7.28 \times 10^{-2}$), p = 0.21 | $\beta = 4.90 \times 10^{-2}$ (SE = $1.34 \times 10^{-2}$), p = $2.6 \times 10^{-4}$ |
| MDD score<br>pT = 0.05 | $\beta = 7.44 \times 10^{-2}$ (95%CI $2.71 \times 10^{-2}$ to $1.18 \times 10^{-1}$), p = $2.1 \times 10^{-3}$ | $\beta = 7.02 \times 10^{-2}$ (SE = $1.35 \times 10^{-2}$), p = $2.1 \times 10^{-7}$ |
| MDD score<br>pT = 0.1 | $\beta = 9.75 \times 10^{-2}$ (95%CI $4.53 \times 10^{-2}$ to $1.35 \times 10^{-1}$), p < $2 \times 10^{-4}$ | $\beta = 7.38 \times 10^{-2}$ (SE = $1.37 \times 10^{-2}$), p = $6.4 \times 10^{-8}$ |
| MDD score<br>pT = 0.5 | $\beta = 9.68 \times 10^{-2}$ (95%CI $4.48 \times 10^{-2}$ to $1.33 \times 10^{-1}$), p < $2 \times 10^{-4}$ | $\beta = 6.11 \times 10^{-2}$ (SE $1.37 \times 10^{-2}$), p = $8.10 \times 10^{-6}$ |

Having established a relationship between polygenic risk of MDD, MDD in GS:SFHS and probable MDD in UK Biobank, we then sought to test the relationship between polygenic risk of MDD and chronic pain. Polygenic risk of MDD was associated with chronic pain in GS:SFHS at three out of four thresholds. The threshold of pT = 0.01 did not show a significant association with chronic pain in GS:SFHS. This was the same threshold that showed no association with MDD in the same study.

In order to confirm the novel association between polygenic risk of MDD and pain in GS:SFHS, we conducted a replication in the UK Biobank sample. 121,052 UKB individuals provided phenotypic and genotypic data for analysis. Polygenic risk of MDD (at 4 thresholds) was positively and significantly associated with each of the four MDD phenotypes in UKB.

**Table 4:** Prediction of chronic pain in Generation Scotland and UK Biobank using polygenic scores for MDD derived from the Psychiatric Genomic Consortium. Beta: represents standardised regression coefficient between the four polygenic risk scores and the two chronic pain phenotypes. All analyses were conducted in MCMCglmm. The 95% CI represents the 95% credible interval, which is broadly interpreted as the interval in which there is a 95% probability that the true parameter lies.

| Polygenic score | Chronic Pain Grade in Generation Scotland | Chronic Pain Grade in UK Biobank |
| --- | --- | --- |
| MDD score<br>$pT = 0.01$ | $\beta = 2.81 \times 10^{-2}$ (95%CI $-5.56 \times 10^{-3}$ to $6.22 \times 10^{-2}$), $p = 9.7 \times 10^{-2}$ | $\beta = 1.56 \times 10^{-2}$ (95%CI $6.36 \times 10^{-3}$ to $2.60 \times 10^{-2}$), $p = 1.5 \times 10^{-3}$ |
| MDD score<br>$pT = 0.05$ | $\beta = 5.49 \times 10^{-2}$ (95%CI $2.40 \times 10^{-2}$ to $9.16 \times 10^{-2}$), $p = 1.70 \times 10^{-3}$ | $\beta = 2.27 \times 10^{-2}$ (95%CI $1.27 \times 10^{-2}$ to $3.21 \times 10^{-2}$), $p < 3 \times 10^{-4}$ |
| MDD score<br>$pT = 0.1$ | $\beta = 6.62 \times 10^{-2}$ (95%CI $2.82 \times 10^{-2}$ to $9.76 \times 10^{-2}$), $p = 4.3 \times 10^{-4}$ | $\beta = 2.56 \times 10^{-2}$ 95%CI $1.62 \times 10^{-2}$ to $3.63 \times 10^{-2}$, $p < 3 \times 10^{-4}$ |
| MDD score<br>$pT = 0.5$ | $\beta = 5.07 \times 10^{-2}$ (95%CI $1.69 \times 10^{-2}$ to $8.35 \times 10^{-2}$), $p = 1.3 \times 10^{-3}$ | $\beta = 2.35 \times 10^{-2}$, 95%CI $1.47 \times 10^{-2}$ to $3.37 \times 10^{-2}$, $p < 3 \times 10^{-4}$ |

**Figure 2:**
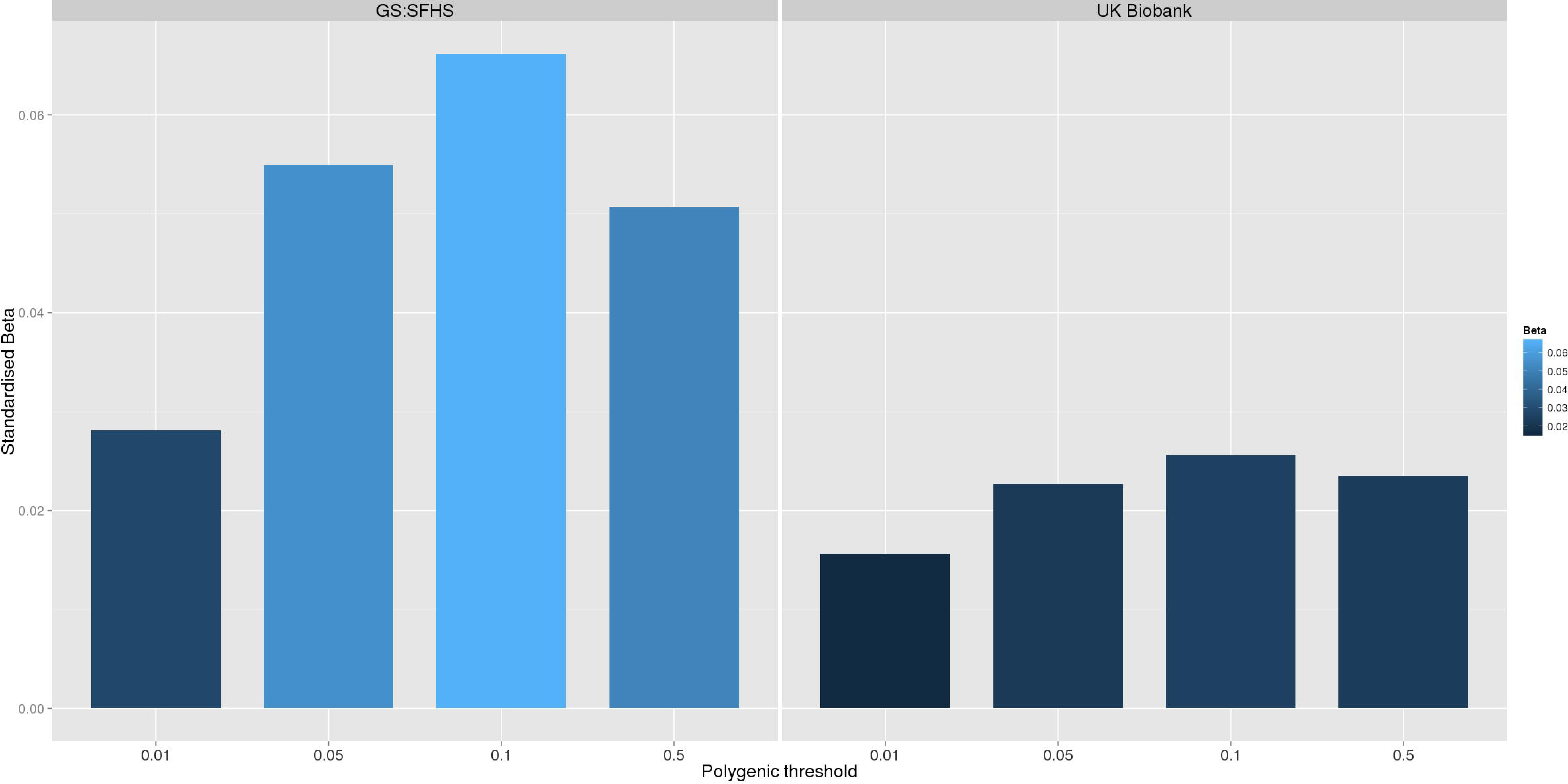
Association between polygenic risk of MDD and chronic pain phenotypes in GS:SFHS and UK Biobank. shows the association between polygenic risk scores for MDD (derived from Psychiatric Genomics Consortium data) and chronic pain in GS:SFHS (left panel) and UK Biobank (right panel). Vertical y-axis represents the effect size as a standardised beta, horizontal axis represents the four alternative p-value thresholds used for the generation of polygenic scores in the discovery GWAS studies.

## Discussion

In the GS:SFHS genetic factors and the environmental effect of spouse/partner contributed significantly to variation in chronic pain defined. Narrow sense heritability of chronic pain was 38.4%, the effect of spouse contributed a further 18.7% to phenotypic variance and a further 42.9% of phenotypic variance was due to residual error, non-shared environment and other unknown sources of variation. Chronic pain was also positively correlated with MDD and this correlation was due to both shared genetic factors and to an effect of spouse/partner.

Using data from two independent GWAS studies (Pfizer-23andMe and PGC), polygenic profile scores for chronic pain predicted both chronic pain traits in GS:SFHS and UK Biobank, but not MDD. In contrast, polygenic risk of MDD was positively and significantly associated with MDD and chronic pain in both GS:SFHS and UK Biobank.

This investigation replicates and extends earlier work showing that chronic pain is a heritable phenotype [28] by demonstrating that there is an independent contribution to risk from the environment shared by an individual’s partner or spouse. We also demonstrate that chronic pain has a polygenic risk architecture, further supporting analyses aimed at identifying specific risk-associated variants. Whilst pain is phenotypically and genetically correlated with MDD, the added contribution of an affected spouse was also substantial in the current study. The reasons for this are unclear in the current study, but the findings imply that the presence of pain in one spouse/partner makes the probability of MDD and higher levels of neuroticism and distress in the other more likely, and vice versa. Possible reasons for this include learned behaviours and the effects of caring for someone with chronic disabling illness. An alternative explanation is that a spousal correlation represents the impact of a common recent environment jointly on both spouses. Spouses are a group currently living together and their correlation could represent common infectious disease history, diet, deprivation, gardening habit and a number of as yet unidentified by shared factors. In addition, an effect of assortative mating cannot be excluded.

Whilst tests of genetic correlation imply shared genetic risk factors, they cannot ascertain the mechanisms of any effect. Similarly, they cannot determine whether the presence of chronic pain in one individual is associated with a risk of depression in a relative more likely because of shared mechanisms or because one phenotype is the consequence of the other. Interestingly, in our analyses, risk of pain was associated with increased genomic risk for MDD. There was, however, no significant association of MDD with genomic risk of pain after correction for multiple testing. These results imply that chronic pain risk (across a number of diverse pathologies) does not strongly predict MDD; whereas polygenic risk of MDD significantly predicts chronic pain. This association may be due to shared mechanisms but it is also possible that a genetic risk of MDD lowers the threshold at which pain is perceived or reported, even if an individual does not develop frank psychiatric disorder. The finding of a robust MDD polygenic risk score association with pain is somewhat unexpected, as MDD has (to date) not been particularly tractable to genetic studies. This finding implies that (against the odds) we have been able to detect a specific and potentially important genetic relationship between MDD and pain.

Several limitations should be considered when interpreting the findings from this study. Whilst GS:SFHS is a large family-based study, the genetic effect estimated in the pedigree based study tend to be inflated by shared environmental factors. Although we applied stepwise model selection to include the environment factor with relatively large effect size in the model, there existed the possibility that environment factors with relatively small effect size could not be accounted for. Whilst the use of the DIC criterion for non-Gaussian models is subject to several important limitations [29], the lower DIC values for the models including spouse only are less dependent on sample size, and thus may provide additional information to the comparison of log-likelihoods. Secondly, the use of polygenic risk scores depends largely on the precision of the marker estimates in the independent discovery set. The precision of these estimates from Pfizer-23andMe and the PGC are dependent upon a number of factors, including sample size, allele frequency and heritability. The lack of an association with polygenic risk of chronic pain and MDD may be due in part to poorly estimated marker effect sizes, or to unidentified factors related to differences in the cohorts. However, in contrast to complex phenotypes of comparable heritability, GWAS studies of MDD in individuals of European ancestry have so far yielded no significant findings. It is perhaps surprising, therefore, that risk of MDD should be significantly associated with chronic pain in both GS:SFHS and UK Biobank. Whilst this finding could be due to pleiotropy, or a directional effect of polygenic risk of MDD on pain-there is a possibility that some cases of MDD are misclassified as chronic pain, leading to inflated estimates of genetic correlation and polygenic association with MDD. Such a misclassification could occur for a number of reasons, including the voicing or manifestation of distress through physical rather than psychological language.

Arguably, one potential limitation to the current study is the use of different definitions of chronic pain and MDD in GS:SFHS and UKB. In spite of this difference, the presence of an association between polygenic risk of MDD and chronic pain in both cohorts, further highlights the strength and robustness of the pain-MDD relationship. The findings also signpost the availability of 3 suitable cohorts (GS:SFHS, UKB and Pfizer-23andMe) of sufficient size and power in which to conduct molecular genetic studies to better understand the shared mechanisms of chronic pain and MDD.

## Acknowledgements

This investigation was supported by Wellcome Trust Grant 104036/Z/14/Z. This research has been conducted using the UK Biobank Resource established using funding from the Wellcome Trust Medical Research Council, Scottish Government and others. Generation Scotland received core support from the Chief Scientist Office of the Scottish Government Health Directorates [CZD/16/6] and the Scottish Funding Council [HR03006]. We would like to thank the research participants and employees of 23andMe for making this work possible.

---

**Supplementary Table 1:** Best fitting models for chronic pain in GS:SFHS.

|  | MCMCglmm Test | ASReml-R LogLik test | Effect sizes (95%CI) |
| --- | --- | --- | --- |
| Additive only (A) | DIC = 31176 | - | A = 35.54% (31.91% to 41.78%) |
| A + sib | DIC = 31087 | $\chi^2 = 4.58$, p = 0.03 | A = 37.04% (31.37% to 42.93%)<br>Sib = 0.06% ($7.26 \times 10^{-3}\%$ to 2.73%) |
| A + spouse | DIC = 28787 | $\chi^2 = 27.23$, p = $1.80 \times 10^{-7}$ | A = 38.29% (33.06% to 44.03%)<br>Spouse = 18.66% (10.83% to 25.06%) |
| A + household | DIC = 30885 | $\chi^2 = -4.06$, p = 0.13 | A = 32.23% (26.17% to 40.00%)<br>Household = 5.65% (2.08% to 11.58%) |
| A + spouse + sib | DIC = 28956 | $\chi^2 = 0.50$, p = 0.48 | A = 37.11% (32.03% to 42.28%)<br>Spouse = 15.83% (9.30% to 23.15%)<br>Sib = 0.20% (0.11% to 2.70%) |
| A + spouse + household | DIC = 28902 | $\chi^2 = -3.17 \times 10^{-5}$, p = 1.0 | A = 38.52% (31.75% to 44.76%)<br>Spouse = 21.15% (8.81% to 25.24%)<br>Household = 0.09% (0.04% to 6.57%) |

DIC: Deviance Information Criterion. Household represents the designated effects from the 'old household' and 'young household'

## References

1. Nicholl BI, Mackay D, Cullen B, Martin DJ, Ul-Haq Z, Mair FS, et al. Chronic multisite pain in major depression and bipolar disorder: cross-sectional study of 149,611 participants in UK Biobank. BMC psychiatry. 2014;14:350. doi:10.1186/s12888-014-0350-4.

2. Diatchenko L, Nackley AG, Tchivileva IE, Shabalina SA, Maixner W. Genetic architecture of human pain perception. Trends Genet. 2007;23(12):605-13. doi:10.1016/j.tig.2007.09.004. PubMed PMID: 18023497.

3. Sullivan PF, Daly MJ, O’Donovan M. Genetic architectures of psychiatric disorders: the emerging picture and its implications. Nature reviews Genetics. 2012;13(8):537-51. doi:10.1038/nrg3240.

4. Hocking LJ, Generation S, Morris AD, Dominiczak AF, Porteous DJ, Smith BH. Heritability of chronic pain in 2195 extended families. European journal of pain (London, England). 2012;16(7):1053-63. doi:10.1002/j.1532-2149.2011.00095.x.

5. Junqueira DR, Ferreira ML, Refshauge K, Maher CG, Hopper JL, Hancock M, et al. Heritability and lifestyle factors in chronic low back pain: results of the Australian twin low back pain study (The AUTBACK study). European journal of pain (London, England). 2014;18(10):1410-8. doi:10.1002/ejp.506.

6. Sullivan PF, Neale MC, Kendler KS. Genetic epidemiology of major depression: review and meta-analysis. The American journal of psychiatry. 2000;157(10):1552-62. PubMed PMID: 11007705.

7. Polderman TJ, Benyamin B, de Leeuw CA, Sullivan PF, van Bochoven A, Visscher PM, et al. Meta-analysis of the heritability of human traits based on fifty years of twin studies. Nat Genet. 2015;47(7):702-9. doi:10.1038/ng.3285. PubMed PMID: 25985137.

8. Sullivan PF, Daly MJ, Ripke S, Lewis CM, Lin DY, Wray NR, et al. A mega-analysis of genome-wide association studies for major depressive disorder. Molecular Psychiatry. 2013;18(4):497-511. doi:10.1038/mp.2012.21. PubMed PMID: WOS:000316568600016.

9. Lubke GH, Hottenga JJ, Walters R, Laurin C, de Geus EJ, Willemsen G, et al. Estimating the Genetic Variance of Major Depressive Disorder Due to All Single Nucleotide Polymorphisms. Biological Psychiatry. 2012. Epub 2012/04/24. doi:10.1016/j.biopsych.2012.03.011. PubMed PMID: 22520966; PubMed Central PMCID: PMC3404250.

10. Richardson R, Westley T, Gariépy G, Austin N, Nandi A. Neighborhood socioeconomic conditions and depression: a systematic review and meta-analysis. Soc Psychiatry Psychiatr Epidemiol. 2015. doi:10.1007/s00127-015-1092-4.

11. Morgan CL, Conway P, Currie CJ. The relationship between self-reported severe pain and measures of socio-economic disadvantage. European journal of pain (London, England). 2011;15(10):1107-11. doi:10.1016/j.ejpain.2011.04.010.

12. Goodson NJ, Smith BH, Hocking LJ, McGilchrist MM, Dominiczak AF, Morris A, et al. Cardiovascular risk factors associated with the metabolic syndrome are more prevalent in people reporting chronic pain: results from a cross-sectional general population study. Pain. 2013;154(9):1595-602. doi:10.1016/j.pain.2013.04.043. PubMed PMID: 23707277.

13. Barnett K, Mercer SW, Norbury M, Watt G, Wyke S, Guthrie B. Epidemiology of multimorbidity and implications for health care, research, and medical education: a cross-sectional study. Lancet. 2012;380(9836):37-43. doi:10.1016/S0140-6736(12)60240-2. PubMed PMID: 22579043.

14. Krein SL, Heisler M, Piette JD, Makki F, Kerr EA. The effect of chronic pain on diabetes patients’ self-management. Diabetes care. 2005;28(1):65-70. PubMed PMID: 15616235.

15. Sudore RL, Karter AJ, Huang ES, Moffet HH, Laiteerapong N, Schenker Y, et al. Symptom burden of adults with type 2 diabetes across the disease course: diabetes & aging study. J Gen Intern Med. 2012;27(12):1674-81. doi:10.1007/s11606-012-2132-3. PubMed PMID: 22854982; PubMed Central PMCID: PMCPMC3509316.

16. Burri A, Ogata S, Vehof J, Williams F. Chronic widespread pain: clinical comorbidities and psychological correlates. Pain. 2015;156(8):1458-64. doi:10.1097/j.pain.0000000000000182. PubMed PMID: 25851458.

17. Smith BH, Campbell A, Linksted P, Fitzpatrick B, Jackson C, Kerr SM, et al. Cohort Profile: Generation Scotland: Scottish Family Health Study (GS:SFHS). The study, its participants and their potential for genetic research on health and illness. International journal of epidemiology. 2013;42(3):689-700. doi:10.1093/ije/dys084.

18. Smith BH, Campbell H, Blackwood D, Connell J, Connor M, Deary IJ, et al. Generation Scotland: the Scottish Family Health Study; a new resource for researching genes and heritability. BMC Med Genet. 2006;7:74. Epub 2006/10/04. doi:1471-2350-7-74 [pii] 10.1186/1471-2350-7-74. PubMed PMID: 17014726.

19. Hocking LJ, Smith BH, Jones GT, Reid DM, Strachan DP, Macfarlane GJ. Genetic variation in the beta2-adrenergic receptor but not catecholamine-O-methyltransferase predisposes to chronic pain: results from the 1958 British Birth Cohort Study. Pain. 2010;149(1):143-51. doi:10.1016/j.pain.2010.01.023. PubMed PMID: 20167428.

20. Pain IAftSo. Classification of chronic pain. Descriptions of chronic pain syndromes and definitions of pain terms. Prepared by the International Association for the Study of Pain, Subcommittee on Taxonomy. Pain Suppl. 1986;3:S1-226. PubMed PMID: 3461421.

21. Purves AM, Penny KI, Munro C, Smith BH, Grimshaw J, Wilson B, et al. Defining chronic pain for epidemiological research. assessing a subjective definition. 1998;10(3):139-47.

22. Von Korff M, Ormel J, Keefe FJ, Dworkin SF. Grading the severity of chronic pain. Pain. 1992;50(2):133-49. PubMed PMID: 1408309.

23. Smith BH, Penny KI, Purves AM, Munro C, Wilson B, Grimshaw J, et al. The Chronic Pain Grade questionnaire: validation and reliability in postal research. Pain. 1997;71(2):141-7. PubMed PMID: 9211475.

24. Fernandez-Pujals AM, Adams MJ, Thomson P, McKechanie AG, Blackwood DH, Smith BH, et al. Epidemiology and Heritability of Major Depressive Disorder, Stratified by Age of Onset, Sex, and Illness Course in Generation Scotland: Scottish Family Health Study (GS:SFHS). PLoS One. 2015;10(11):e0142197. doi:10.1371/journal.pone.0142197. PubMed PMID: 26571028; PubMed Central PMCID: PMCPMC4646689.

25. Allen NE, Sudlow C, Peakman T, Collins R, Biobank UK. UK biobank data: come and get it. Sci Transl Med. 2014;6(224):224ed4. doi:10.1126/scitranslmed.3008601. PubMed PMID: 24553384.

26. Gilmour AR, Gogel BJ, Cullis BR, Thompson R. ASReml user guide release 3.0. VSN International Ltd. 2009.

27. Nakagawa S, Schielzeth H. A general and simple method for obtaining R2 from generalized linear mixed-effects models. Methods in Ecology and Evolution. 2013;4(2):133-42. doi:10.1111/j.2041-210x.2012.00261.x.

28. Hocking LJ, Generation S, Morris AD, Dominiczak AF, Porteous DJ, Smith BH. Heritability of chronic pain in 2195 extended families. European journal of pain. 2012;16(7):1053-63. doi:10.1002/j.1532-2149.2011.00095.x. PubMed PMID: 22337623.

29. Spiegelhalter DJ, Best NG, Carlin BP, Van Der Linde A. Bayesian measures of model complexity and fit. Journal of the Royal Statistical Society: Series B (Statistical Methodology). 2002;64(4):583-639. doi:10.1111/1467-9868.00353.

